# Inferring plant community phenology via bee-collected pollen

**DOI:** 10.1101/2024.08.23.609319

**Authors:** Sydney B. Wizenberg, Mateus Pepinelli, Bao Ngoc Do, Mashaba Moubony, Darya Tamashekan, Ida M. Conflitti, Amro Zayed

**Author notes:** **Author contributions:** AZ and IMC provided supervision and project management. AZ and SBW designed the experiment. SBW, MP, BND, MM, and DT performed the experiment. SBW analyzed the data and wrote the manuscript. All authors revised the manuscript.

## Abstract

Global climate change is producing novel biospheric conditions, presenting a threat to the stability of ecological systems and the health of the organisms that reside within them. Variation in climatic conditions is expected to facilitate phenological reshuffling within plant communities, impacting the plant-pollinator interface, and the release of allergenic pollen into the atmosphere. Impacts on plant, invertebrate, and human health remain unclear largely due to the variable nature of phenological reshuffling and insufficient monitoring of these trends. Large-scale temporal surveillance of plant community flowering has been difficult in the past due to logistical constraints. To address this, we set out to test if metabarcoding of honey bee collected pollen could be used to infer the phenology of plant communities via comparison to *in situ* field monitoring. We found that honey bees can accurately indicate the onset of anthesis, but not its duration, in the plant species they selectively forage on. Increasing the number of colonies used to monitor, and employing a multi-locus approach for metabarcoding of pollen, substantially increased the species detection power of our approach. Here, we demonstrate that metabarcoding of honey bee collected pollen can substantively streamline the establishment of long-term phenological monitoring programs to document the on-going consequences of global climate change and its impact on the temporal aspects of plant-pollinator relationships.

## INTRODUCTION

Plant flowering phenology, e.g. the onset and duration of blooming, is an important life-history trait that guides patterns of reproductive success (Rodríguez-Pérez & Traveset, 2016; Munguía-Rosas et al., 2011; Cooper et al., 2011), interactions with pollinators (Gallagher & Campbell, 2020; Rafferty et al., 2011), and the influx of allergenic pollen into the atmosphere (Ziska & Beggs, 2012). Climate change is expected to broadly augment patterns of annual temperature and rainfall (Allan et al., 2020; Parmesan, 2006), shifting the regionality of climate zones (Anderson et al., 2020), and atmospheric feedback cycles (Xie et al., 2015). This will induce shifts to the spatial-temporal dynamics around plant flowering phenology (Tun et al., 2021; Hamann et al., 2021), presenting a catalyst for global ecological change via reshuffling of community dynamics and plant-pollinator networks. Monitoring the scope and impact of these changes across multiple geographic gradients is incredibly valuable, but traditional techniques, e.g. ground-based historical measurements (Fitchett et al., 2015), are laborious and limited in their scope. The recent popularity of metagenetic work has demonstrated its value for quantifying biological communities (Keck et al., 2022; Gold et al., 2021; Littlefair et al., 2019; Lacoursière-Roussel et al., 2018; Deiner et al., 2017), and could present a new opportunity to modernize this field and facilitate large-scale, standardized monitoring of plant community phenology.

Flowering behaviour in plants is phenotypic; though the abiotic triggers that determine the onset of flowering (e.g. temperature, photoperiodism) are genetically determined (Qanmber et al., 2019; Wellmer and Riechmann, 2010), the actual arrival of those triggers depends on the local environment (Franks et al., 2014). As a result of this, climate change and its impact on regional weather conditions is expected to shift the on-set and duration of flowering in many species, inducing reshuffling of community-level flowering structure (Rafferty et al., 2020). This will drive broad evolutionary and ecological changes (Toju et al., 2017) as it will impact plant populations in a myriad of ways, including their demographic structure (Rhymer et al., 2010; Weis et al., 2005; Weis and Kossler, 2004). Reshuffling of flowering behaviour, at both the population and community level, is expected to impact a variety of evolutionary mechanisms (Bonner et al., 2019; Wadgymar and Weis, 2017; Rymer et al., 2010) and dynamics (Lü et al., 2023; Savage, 2019; Franks and Weis, 2008). This presents a formidable threat to plant communities, as species with high adaptive potential may quickly respond (Franks et al., 2007), while others may struggle to survive under these new conditions. Due to the phenotypic nature of flowering phenology, and the variable nature of phenological reshuffling, the context around these changes and how individual species respond will vary across spatial-temporal gradients (Rafferty et al., 2020; Shen et al., 2015), meaning that empirical work undertaken with a narrow scope is unlikely to provide meaningful information about community-level effects.

Because flowering phenology is a major determinant of plant-pollinator interactions, it’s reasonable to assume that phenological reshuffling will coincide with changes to the temporal aspect of these interactions. In plants, selection on flowering characteristics is often modulated by biotic interactions (Elzinga et al., 2007), and thus any changes to the context around plant-pollinator interactions could have evolutionary and ecological consequences. This is further exacerbated by potential changes to the developmental timing of pollinating insects; reshuffling of insect phenology may differ from that of the plants they interact with. Mismatched synchronicity within plant-pollinator networks could reduce the availability of mutualistic partners, leading to demographic consequences (Hegland et al., 2009), and reductions in population-level fitness (Morton and Rafferty, 2017). This may induce dispersal limitations in entomophilous plants (Vasiliev and Greenwood, 2021) and broadly change the evolutionary ecology of insect-mediated pollination (Inouye, 2020). Temporal reshuffling of flowering phenology thereby represents a potent evolutionary and ecological pressure to both plant communities and the invertebrates that rely on them – monitoring these changes and how they vary across space and time is a crucial step to understanding the far-reaching effects of climate change and its impact on terrestrial ecosystems.

Traditional approaches that rely on observational data have highlighted the value of understanding spatial-temporal flowering dynamics (Rafferty et al., 2020), but have underscored the logistical constraints of monitoring this on a large scale. Current monitoring programs typically entail physical documentation of flowering behaviour at a study site, which is both time consuming, spatially limited, and hindered by its reliance on morphological species identification. Implementing a modern, high-throughput approach could revolutionize this field, as seen when eDNA techniques were applied to monitor biodiversity more broadly (Ruppert et al., 2019). Pollen metabarcoding, a contemporary method for rapidly characterizing the diversity of pollen collected by invertebrates (Bell et al., 2022), may be the key to exploring climate-driven changes to plant phenology. Characterizing plant community structure via metabarcoding of air samples is possible (Johnson et al., 2021) but limited in its ability to identifying phenological patterns due to the presence of non-reproductive vascular plant tissue in filtration systems (Johnson et al., 2019). Collecting reproductive tissue, e.g. pollen, from plants, refines the utility of this approach to limit detection to only species which were blooming during the survey period. Though air-filtration systems would be incapable of discriminating between types of plant tissue, assistance from generalist pollinators could present an opportunity to quickly characterize large populations of flowering plants (Bell et al., 2016). The western honey bee, *Apis mellifera*, is a prolific generalist pollinator (Quigley et al., 2019; Potts et al., 2010), making them ideal candidates for rapidly collecting large quantities of pollen from study sites. Metabarcoding of honey bee collected pollen to monitor temporal reshuffling of plant communities could revolutionize our ability to document phenological changes on a previously unfathomable scale. However, it currently remains unclear if pollen collected by honey bees reliably predicts the phenology of flowering plants. To address this knowledge gap, we set out to test the reliability of honey bee assisted phenological monitoring via comparison to in situ field measurements, and explore the boundaries of this approach.

## MATERIALS AND METHODS

### Experimental design

We used a rooftop apiary located on York University’s Keele campus as our focal study site (Fig. 1). York University is located in Toronto, Ontario, situated along the northern border of the Carolinian region. This ecologically rich area is characterized as a biodiversity hotbed as its estimated to host over 2300 plant species (Almas and Conway, 2016), some of which are endemic to this region. At the beginning of the beekeeping season, we randomly selected 5 colonies for use in our monitoring program. We installed pollen traps (Pollen Trap Bottom Board, model #APH1000, ApiHex Beekeeping Supplies, Guelph, ON, Canada) at the bottom of each colony and disabled the traps until the commencement of the experiment. At the beginning of May we began our initial 4-week experiment. We used a 1000 meter radius around our rooftop apiary as our field site (Fig. 1), and split the resulting area into 4 equally sized quadrants. For each quadrant, we completed vertical transects, inventorying all species in bloom. All four quadrants were surveyed weekly, and flowering onset was determined by the presence of at least 1 visible set of dehiscent anthers. Pollen collection from the honey bee colonies occurred weekly, using the follow methods: at the start of the day (approximately 9:00am) we blocked off the main hive entrance and enabled the pollen trap entrance. Foraging bees entered the colony via the trap entrance, wherein the pollen clumps were scraped off of their legs by trap mechanisms. The pollen was collected in a metal tray located at the base of the hive. After 24 hours the trap was disabled (and the main hive entrance was re-opened), and the resulting pollen was transferred to a sterile conical tube, weighed in a sterile weigh boat, and transferred to a -20 °C freezer. Field work was discontinued after 4 weeks, but pollen collection continued on a weekly schedule for a total of 15 weeks, encompassing May – August. Samples were stored in the freezer until undergoing pollen metabarcoding at the end of the monitoring program.

**Figure 1:**
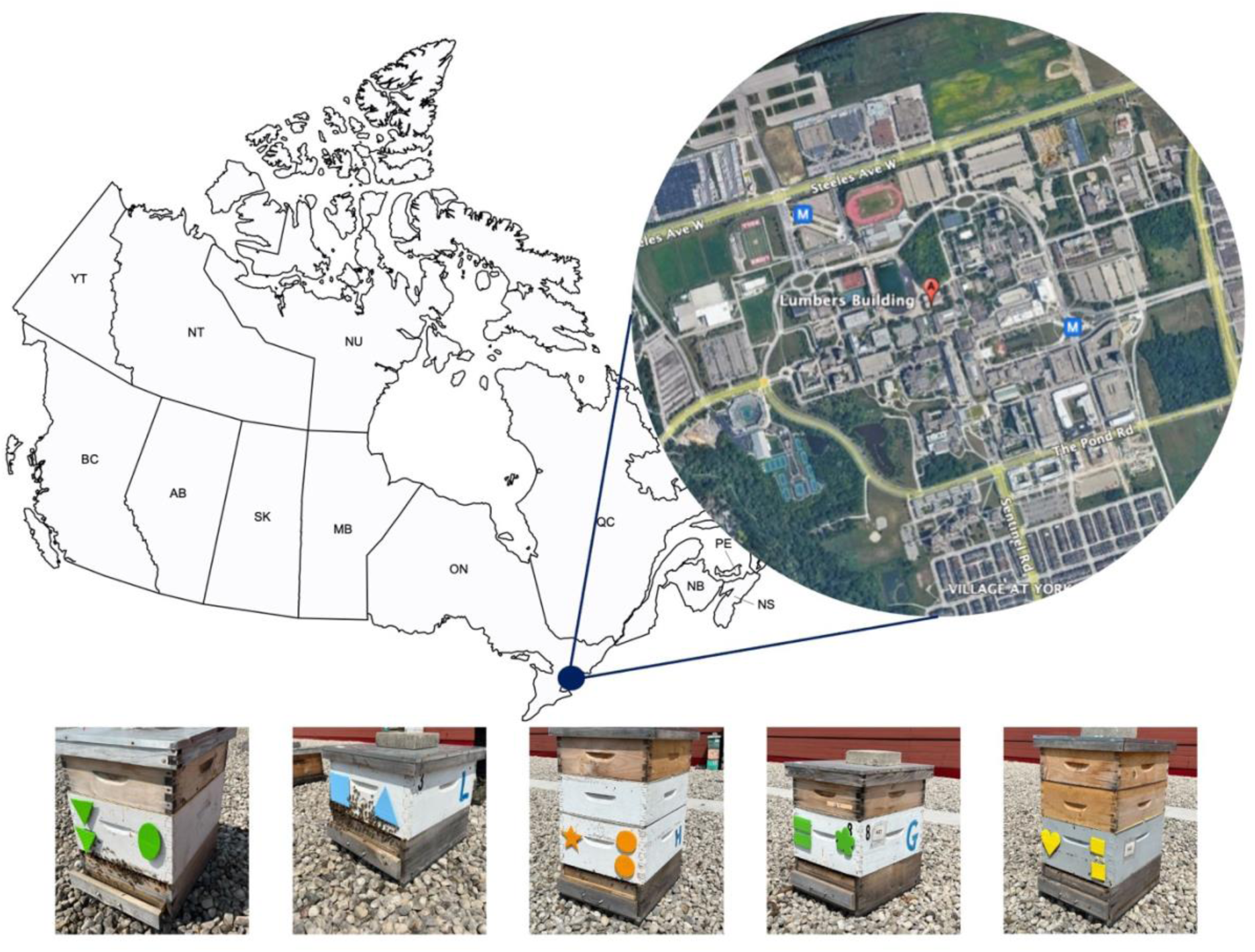
Map of the location of our study site. Our urban apiary is located on the top of the Lumbers building within York University’s campus in Toronto, Ontario, Canada. In situ field monitoring covered the 1000 m. radius surrounding our urban apiary.

### Pollen metabarcoding

We followed an established protocol for multi-locus pollen metabarcoding (Wizenberg et al., 2023a). We extracted DNA from pollen samples using the NucleoMag DNA Food Kit (Macherey-Nagel, Düren, Germany). We combined each bee-collected pollen sample with lysis buffer based on the weighted mass of the sample; for each 1 g. of pollen, we used 2 mL of lysis buffer: 70% autoclaved filtered water (Millipore Sigma, Burlington MA, USA), 20% 10x STE (100 mM NaCl, 10 mM Tris, 25 mM EDTA), and 10% diluted SDS (10% sodium dodecyl sulfate). We then sealed each conical tube, inverted them 10 times, then homogenized the suspension by shaking samples for 10 minutes in an orbital shaker (G25 Incubator Shaker, New Brunswick Scientific, Edison, NJ, USA) at 25°C and 375 rpm. Immediately after removing the samples from the orbital shaker, we pipetted 3 mL of the homogenized sample into a 7 mL cylindrical tube containing 10 small (1.4 mm) and 2 large (2.8 mm) ceramic beads and bead beat it (Bead Mill 24, Fisherbrand, Ottawa, Ont., Canada) for 4 x 30 s cycles, at a speed of 6 m/s. We then transferred 550 uL of the homogenized suspension to a 1.5 mL Eppendorf tube. We warmed CF lysis buffer for 10 min in a 65°C water bath then added 550 uL of the warmed buffer and 10 uL of Proteinase K (from the NucleoMag DNA Food kit) to the homogenized sample and vortexed (Mini Vortex Mixer, VWR, Mississauga, Ont., Canada) the sealed tube for 30 s. We then incubated the sample at 65 °C for 30 min in a block heater (Isotemp 145D, 250V, Fisherbrand, Ottawa, Ont., Canada), inverting every 10 min. After incubation, we added 20 uL of RNase A (New England Biolabs, Ipswich, MA, USA) and allowed the sample to incubate at room temperature (20°C) for an additional 30 min. After incubation, we centrifuged the sample for 20 min at 14,000 rpm (Centrifuge 5810 R, 15 amps, Eppendorf, Hamburg, Germany), transferred 400 uL of the upper liquid layer to the binding plate, added 25 uL of NucleoMag B-Beads and 600 uL of binding buffer CB (both from the NucleoMag DNA Food kit) then ran an extraction program on the KingFisher Flex extraction robot (ThermoScientific, Waltham, MA, USA). Each of the 5 deep well plates used to complete the extraction program contained either 600 uL of CMW buffer (wash 1), 600 uL of CQW buffer (wash 2.1), 600 uL of 80% EtOH (wash 2.2), or 100 uL of buffer CE (elution). After the extraction program was complete, we transferred 80 uL of the eluted sample to a fresh 1.5 mL Eppendorf tube and froze it at -20 °C until we began DNA amplification.

We carried out three PCR programs – first amplifying the DNA barcode locus of interest, then extending the length of the amplified sequence, and finally dual-indexing the samples with unique combinations of forward and reverse primers. We used validated primers (Wizenberg et al., 2023) to amplify loci from two barcoding regions, ITS2 and rbcL1 (Table 1). We used 96 well plates containing 86 pollen samples and 10 controls. Each reaction included 11 uL of water, 12.5 uL of 2x Taq Pol Mix (New England Biolabs, Ipswich, MA, USA), 0.5 uL of each relevant forward and reverse primer (1 µM), and 0.5 uL of sample DNA (∼ 36.2 ng) into each well. PCR cycling conditions were (Eppendorf Mastercycler, Ep Gradient, Hamberg, Germany): initial denaturation (94°C, 10 min, 1 cycle), followed by 40 cycles of denaturation (94°C, 30 s), annealing (54°C, 40 s), and extension (72°C, 1 min), then a final extension cycle (72 °C, 10 min). The product of this first PCR (PCR1) was used as the template for a second PCR reaction (PCR2; Table 1) and the same chemistry as described above (with 0.5 uL of PCR1 product instead of sample DNA). Cycling conditions were the same for PCR1 and PCR2, with the exception of a higher annealing temperature (56°C instead of 54°C). After each respective PCR program, we used gel electrophoresis to confirm sufficient amplification of each sample and identify any potential contamination using the negative controls. Following PCR2, we prepared samples for Illumina Sequencing by performing a third PCR program that tagged each sample with a unique combination of forward and reverse primers; PCR3 program specifications follow that described above, with the exception of a higher annealing temperature of 60°C. We then normalized the resulting PCR3 product using a SequalPrep Normalization kit (Invitrogen, Burlington, ON, Canada), and shipped the normalized libraries on dry ice for Illumina Sequencing (Illumina MiSeq PE250) at Genome Quebec. Each library was pair-end sequenced in its own lane.

**Table 1:**
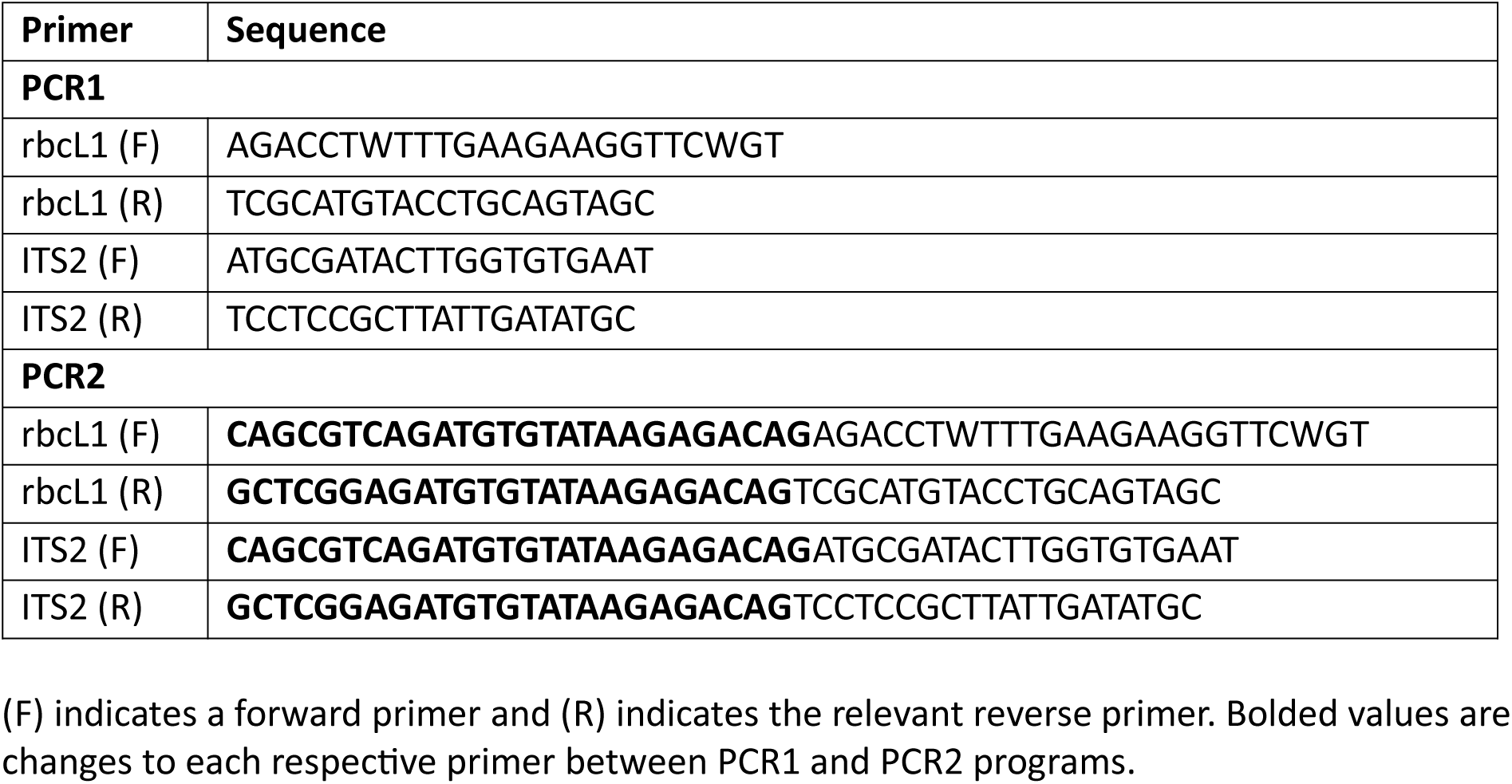
PCR primer specifications.

### Data analysis

Pollen metabarcoding data processing was completed in Python (v. 3.10.7), and R (v. 4.2.1; R Core Team), using the *dada2* (Callahan et al., 2016) (v. 1.16.0) and *purrr* (Henry & Wickham, 2020) (v. 0.3.4) packages. We first paired forward and reverse reads, trimmed primer sequences, and grouped identical sequences as ASVs (amplicon sequence variants). We then built a database that linked species to sequences associated with each primer using the MetaCurator method (Richardson et al., 2020). We used this database to parse through returned sequence data and identify the species associated with each unique grouped sequence, setting a precursory condition of >0.95 similarity. After identifying the plant species associated with each sequence, we consolidated classifications at the genera level, and filtered data to control for sequence mistagging (Richardson, 2022). We used Richardson’s (2022) established filtering method; negative controls acted as indicators of mistagging frequency, and we filtered real sample data to remove detections with a high likelihood of representing mistag-associated false detections (Richardson, 2022). All analysis was completed in R (v. 3.10.7, R Core Team). For correlation tests, we used the cor.test() function, and for analysis of variance we used the aov() function, both included in the base *stats* package (v. 4.2.1; R Core Team). All data visualization was competed using the ggplot() function from *ggplot2* (v. 3.4.2; Wickham, 2016), and the geom_density_ridges() function from the *ggridges* extension (v. 0.5.4; Wilke, 2018).

## RESULTS

### Method validation

Our inventory of flowering plants yielded 51 unique genera, while our temporally paired pollen metabarcoding libraries yielded 99 unique genera. The additional genera detected via pollen metabarcoding indicates that the bees were likely foraging beyond the 1000 meter radius used as our study site. While it remains possible that some of these may be false positives, our method validation suggests that this is unlikely (see species accumulation curves, Wizenberg et al., 2023a), though we do recommend that any researchers undertaking monitoring via this approach use their best judgement when determining if a genera is likely to be present near their study site. Genera suspected to represent a false-positive barcode can be excluded from analysis of phenological structure. Species present in the field but not detected via pollen metabarcoding were either not foraged on by our colonies or may not be suitable for metabarcoding, either as a result of PCR bias or insufficient reference sequence availability.

Species that were present in both datasets (n = 34) were selected to undergo correlation analysis to determine if the onset, duration, and end of flowering could be inferred from pollen metabarcoding data. The onset of flowering in the field was strongly positively associated with the initial occurrence of pollen in our metagenetic datasets (t ∼ ∞, r = 1.0, p < 0.001), for all 34 genera, there was a 1-to-1 rate of agreement among the two approaches. Both the length (t = 7.710, r = 0.806, p < 0.001) and end (t = 5.318, r = 0.685, p < 0.001) of flowering were significantly positively correlated across the two approaches, but did show some degree of disagreement within some of the genera (Fig. 2). Species detection rates differed significantly based on the number of colonies used (F_4,78_ = 175.367, p < 0.001), the genetic approach (F_2,78_ = 750.105, p < 0.001), and their interaction (F_8,78_ = 5.411, p < 0.001). Species detection rates were highest when employing a multi-locus approach with 5 study colonies (Fig. 3). For single-locus characterization, rbcL out-performed ITS2 (Table 2), and across all methods, increasing the number of study colonies directly increased the equivalent species detection rates (Fig. 3).

**Figure 2:**
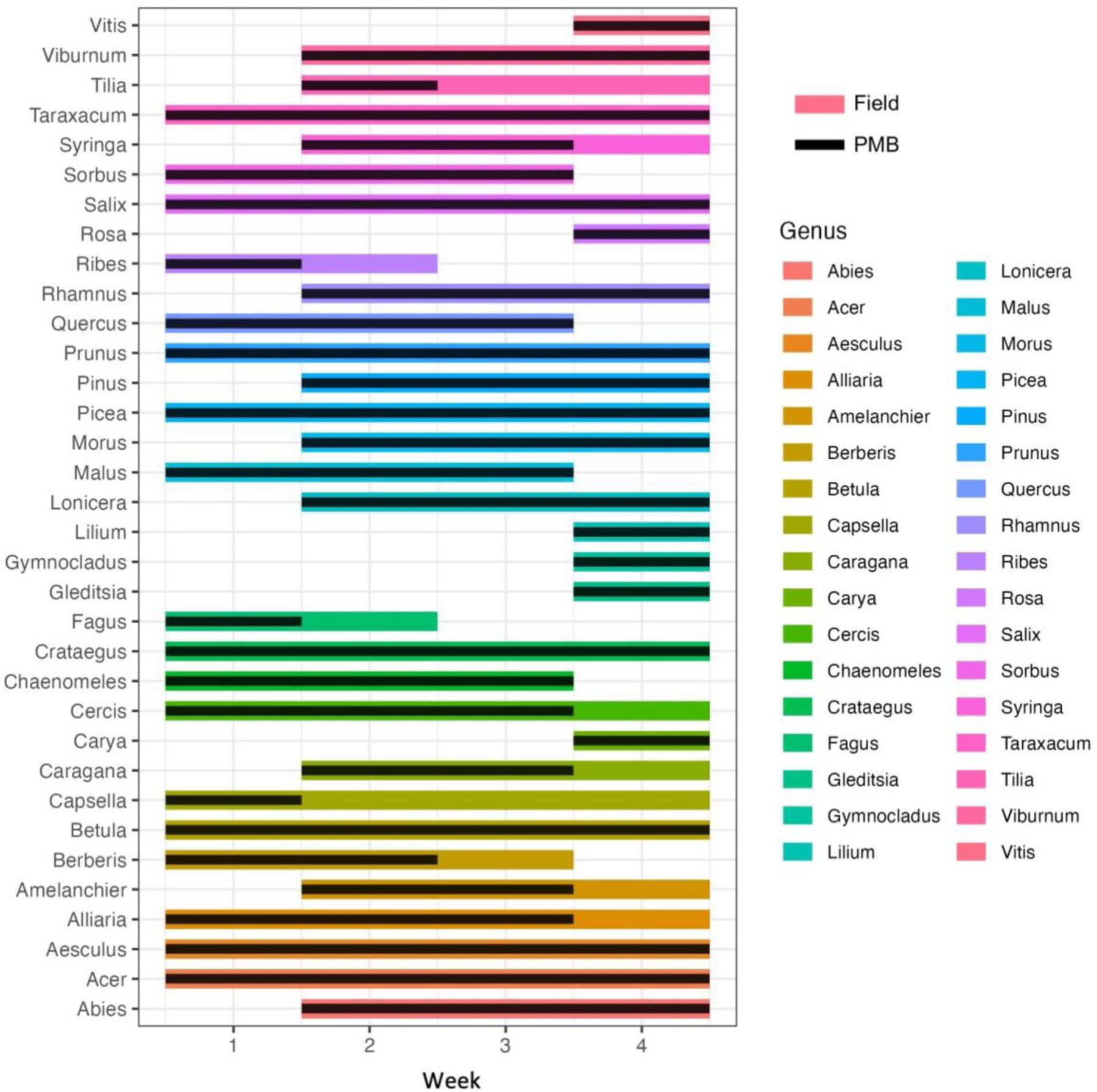
Temporally paired comparisons of pollen metabarcoding data (PMB) and in situ field observations (Field). 34 genera were present in both datasets and underwent correlation analysis to determine if pollen metabarcoding data could be used to infer phenological behaviour. The start, duration, and end of flowering were all significantly positively correlated across the two methods (p < 0.05).

**Figure 3:**
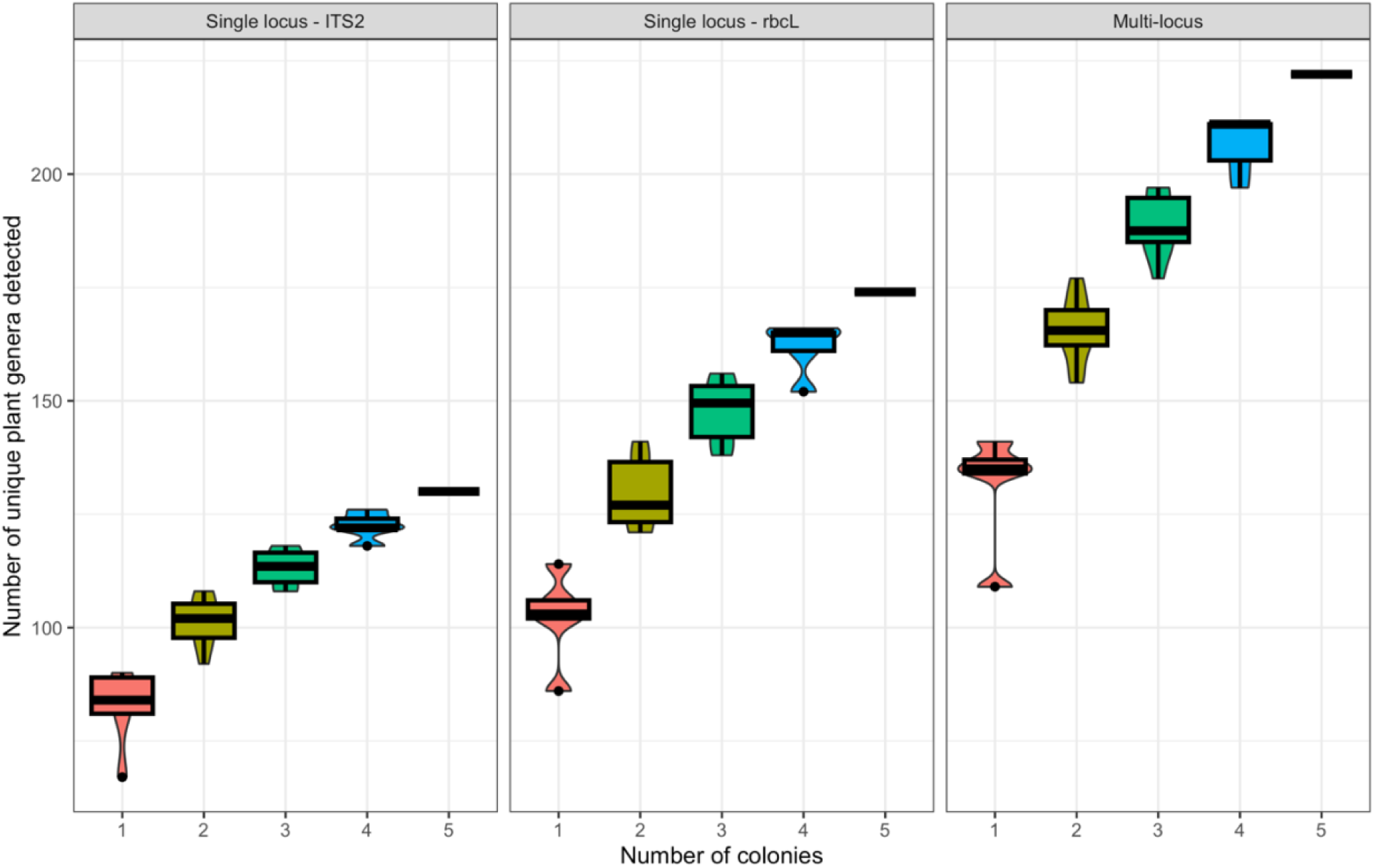
Violin boxplots of unique genera detection rates based on the number of colonies and the loci used for monitoring. Detection rates differed significantly based on the number of colonies used (F_4,78_ = 175.367, p < 0.001), the genetic approach (F_2,78_ = 750.105, p < 0.001), and their interaction (F_8,78_ = 5.411, p < 0.001).

**Table 2:** Species detection rates.

| # of colonies | Single locus – ITS2 |  | Single locus - rbclL |  | Multi-locus |  |
| --- | --- | --- | --- | --- | --- | --- |
|  | Average | SD | Average | SD | Average | SD |
| 1 | 82.2 | 9.26 | 102.2 | 10.21 | 131.2 | 12.7 |
| 2 | 101.4 | 5.27 | 129.4 | 7.47 | 165.5 | 7.03 |
| 3 | 113.3 | 3.89 | 147.6 | 6.79 | 188.5 | 6.74 |
| 4 | 122.4 | 2.97 | 161.8 | 5.81 | 206.6 | 6.39 |
| 5 | 130 | -- | 174 | -- | 222 | -- |

Average is the number of unique genera detected based on the number of colonies used to monitor and the metagenetic approach used for analysis. SD is the standard deviation associated with each average.

### Temporal monitoring

Across our 15-week monitoring program, DNA metabarcoding of honey bee collected pollen from our five colonies yielded the identification of 222 unique plant genera (Fig. 4; interactive version: Appendix 1). Across all 5 colonies, 24-hr. foraging yield was variable but significantly decreased throughout time (Fig. 5). One of our colonies (#2) ceased collecting usable quantities of pollen at the end of June; among the 5 colonies used in our study, this colony consistently collected the lowest quantities for the duration of the monitoring program. Three of the five colonies consistently provided usable quantities of pollen (> 1 g.) across the entire duration of our monitoring program (Fig. 6). Among those three colonies, dietary composition varied, but dominant dietary sources of pollen were relatively consistent (Fig. 6), with substantial contributions from Asteraceae, Fabaceae, and Rosaceae. Foraging yield was significantly correlated with species diversity (t = 2.484, r = 0.296, p = 0.016) but not richness (t = 1.1, r = 0.136, p = 0.275); increases in the amount of pollen collected during a 24-hr. period was associated with greater dietary diversity (Fig. 7b), but not richness (Fig. 7a). Across time, dietary richness (t = -1.066, r = -0.250, p = 0.043) and diversity (t = -2.126, r = -0.257, p = 0.037) significantly declined (Fig. 7c,d). Taxonomic grouping of flowering genera density showed preliminary evidence of phylogenetic clustering at both the family and order level (Fig. 8).

**Figure 4:**
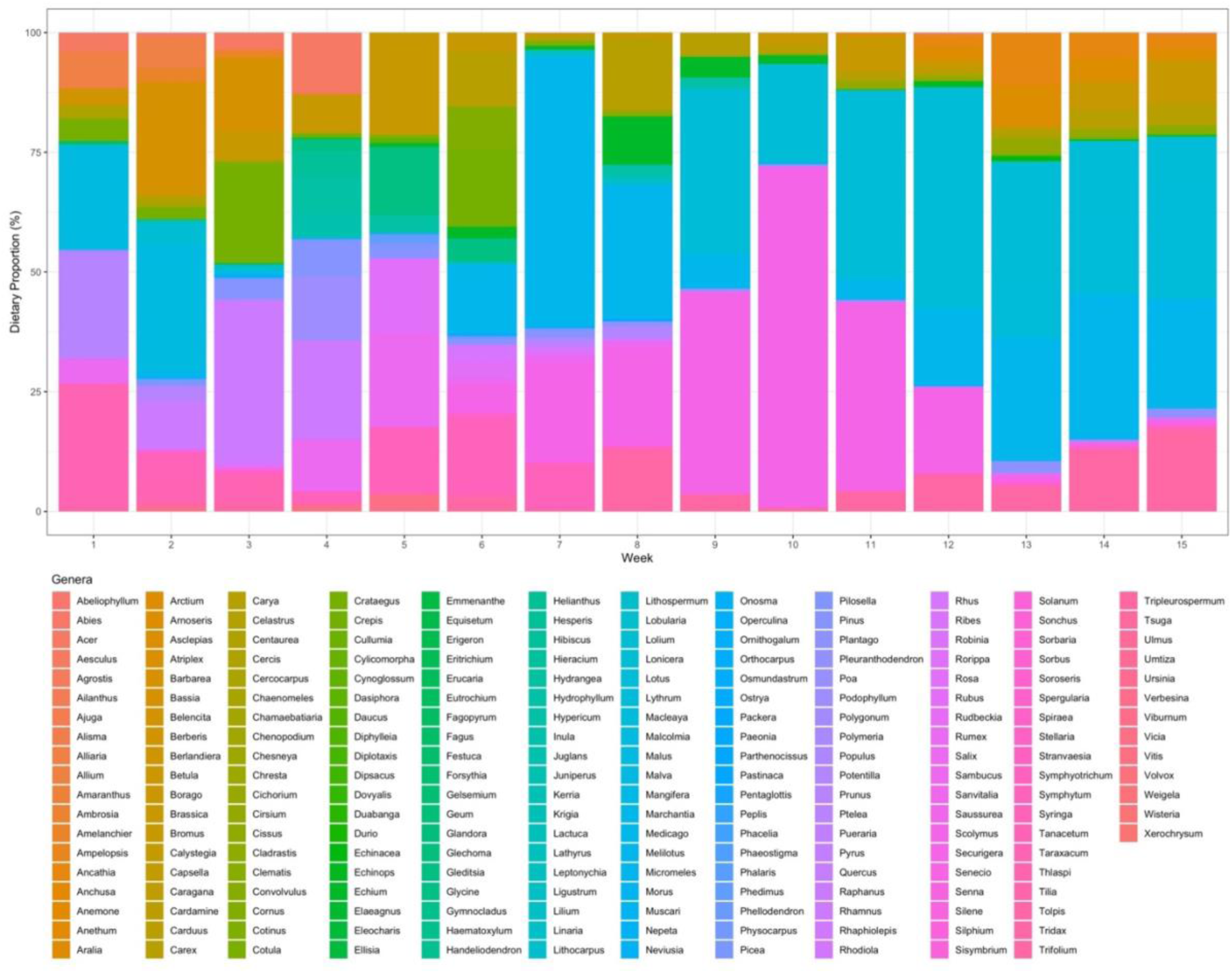
Dietary composition of the 5 honey bee colonies at our urban monitoring site. Colours represent the relative abundance of reads associated with each of the 222 plant genera detected in our pollen samples, estimated via multi-locus pollen metabarcoding. Note: an interactive version of this figure is available as a supplementary file (Appendix 1).

**Figure 5:**
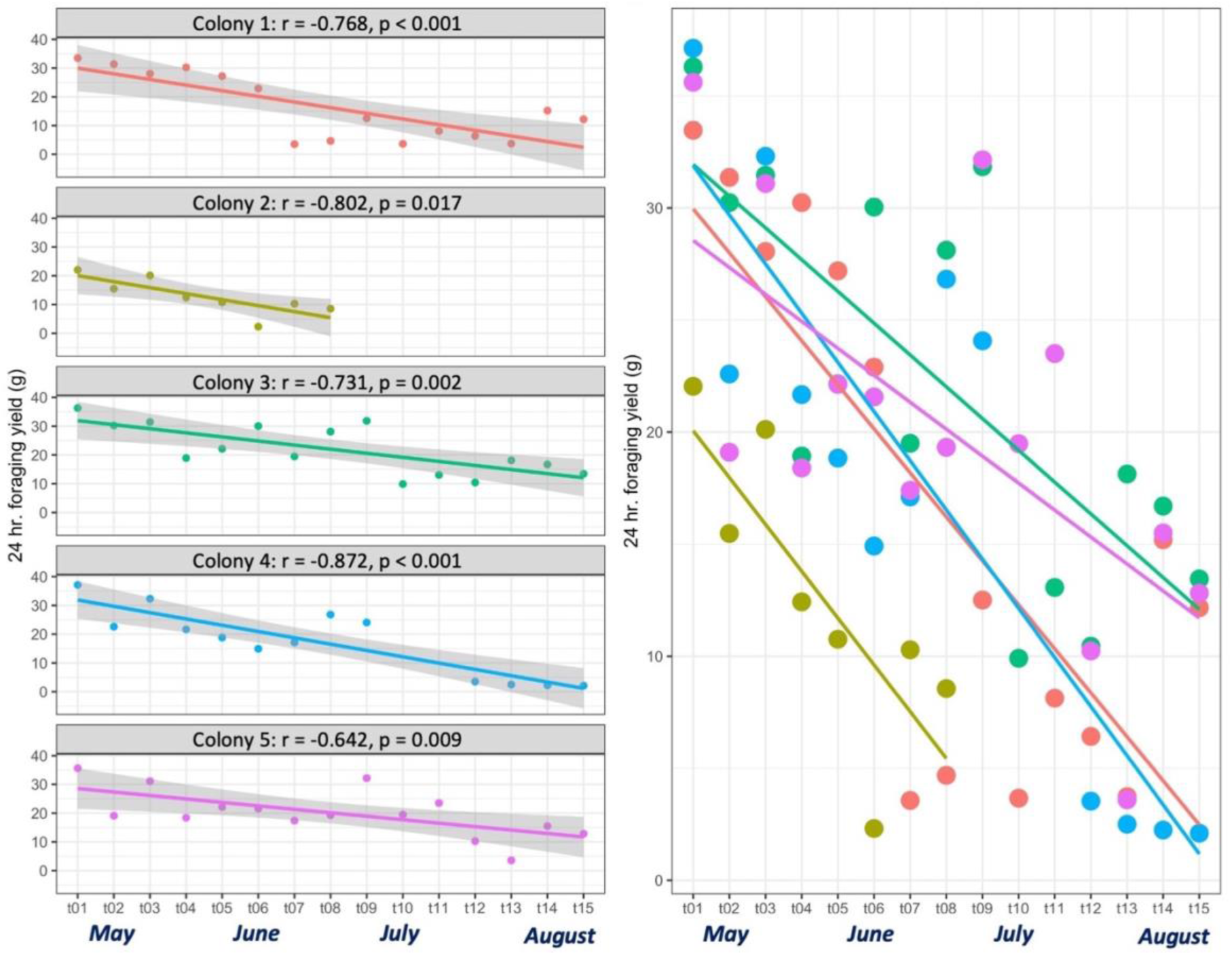
Declines in the 24 hour foraging yield of our urban honey bees across time. Total pollen yield declined significantly across time (p < 0.05) for all 5 of our study colonies. One colony (#2) failed to provide usable pollen yields after the 8th week of monitoring.

**Figure 6:**
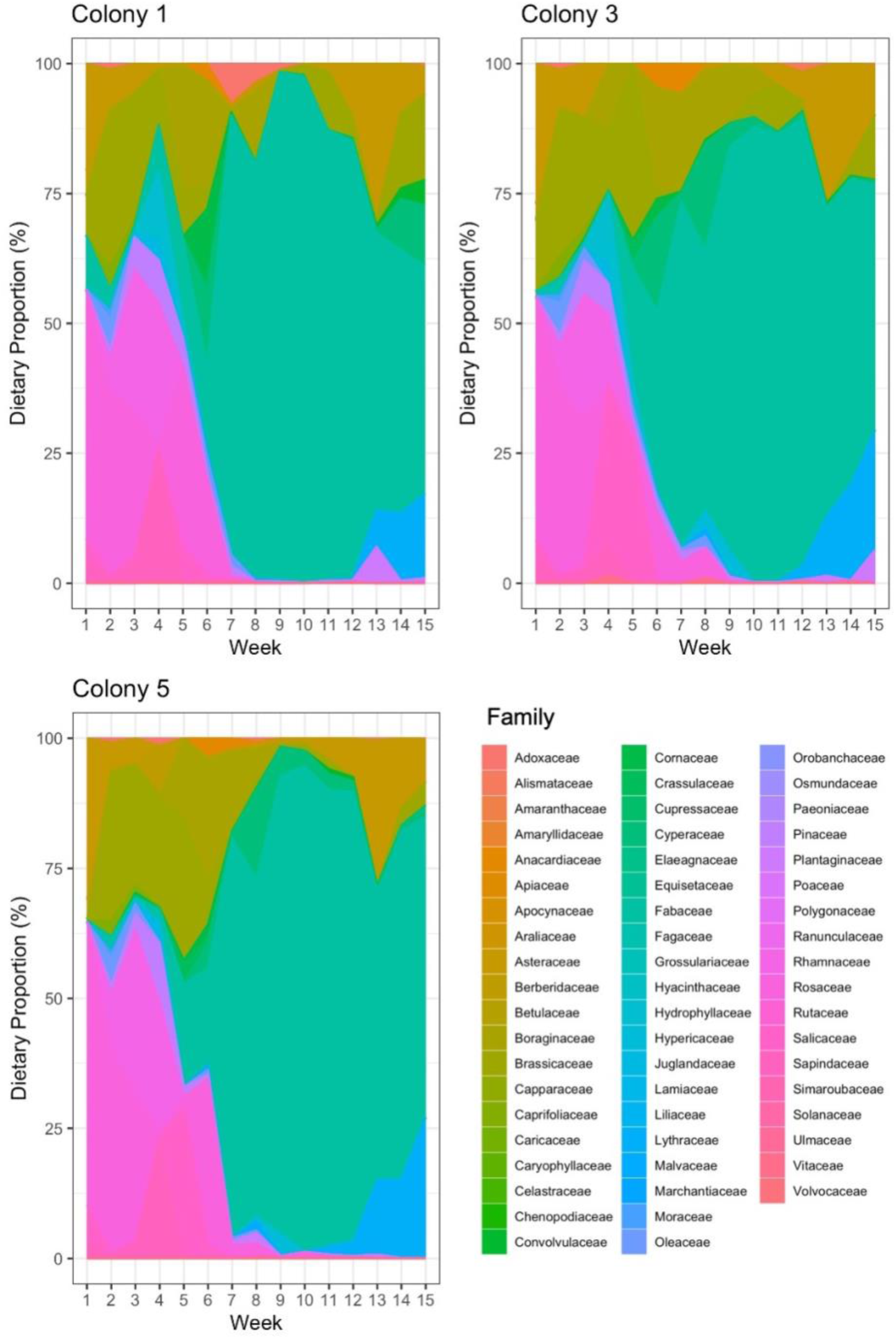
Family-level dietary composition of the 3 colonies with complete time series. Across all three colonies that provided usable pollen samples for the duration of the 15-week monitoring program, the same major plant families were dominant in their diets.

**Figure 7:**
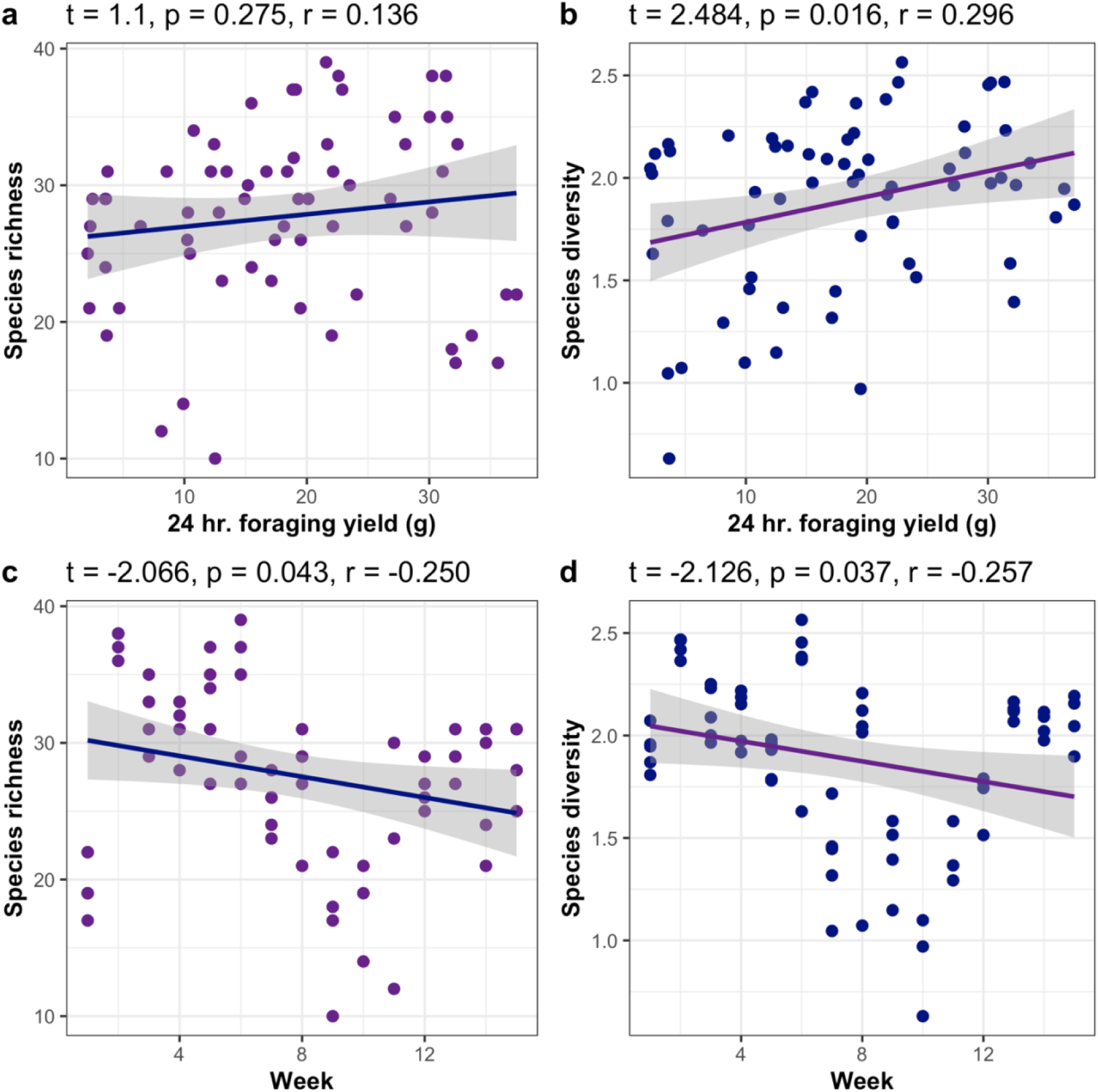
Species richness (p = 0.043) and diversity (p = 0.037) significantly declined across the duration of our monitoring program. Diversity was significantly negatively associated with 24 hr. foraging yield (p = 0.016), but richness was not (p = 0.275).

**Figure 8:**
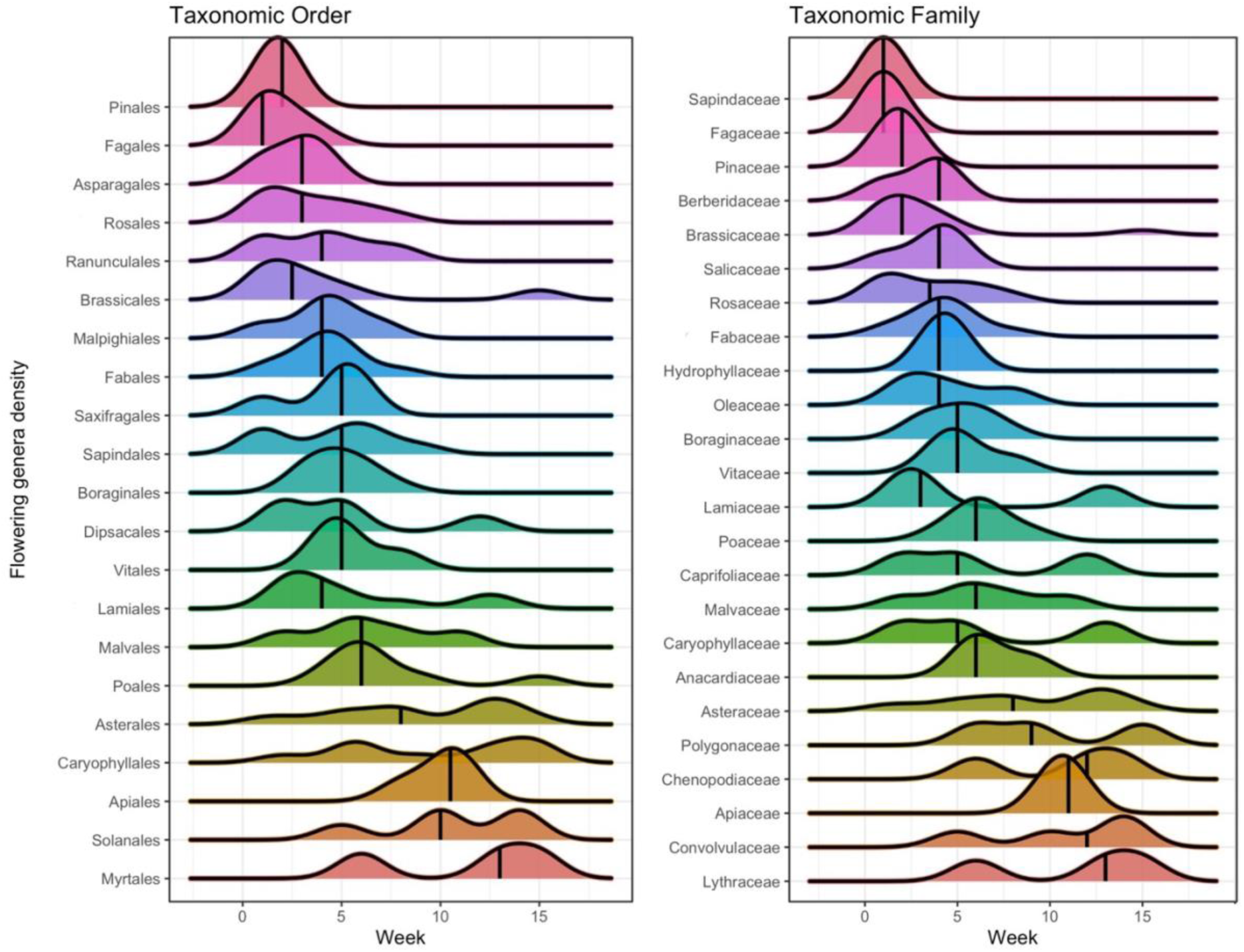
Ridge plots showing temporal clustering of flowering behaviour at upper taxonomic levels. Flowering genera density is the number of genera from that taxonomic group that began flowering during that week, detected via multi-locus metabarcoding of honey bee collected pollen.

## DISCUSSION

Our work has demonstrated that metabarcoding of honey bee collected pollen can be used to infer the phenological structure of flowering plant communities, with some reasonable limitations. When comparing in situ field observations to temporally-paired metabarcoded pollen samples, we found that the onset of flowering could be precisely inferred (Fig. 2). Both the length and end of flowering, though significantly positively correlated among the two methods, showed a greater degree of variation (Fig. 2) and thus may be unreliable indicators of real-life flowering behaviour. This is not surprising, as honey bees are known to rapidly discover and abandon floral resources (Tereshko & Lee, 2002). Foraging bees that travel outside of the colony may sample a large number of pollen sources, but mass pollination events typically only occur after locating favorable dietary sources (Tereshko & Lee, 2002), indicated via a waggle dance (Shackleton et al., 2023; Richardson et al., 2019; Ratnieks & Shackleton, 2015; Biesmeijer & Seeley, 2005). Thus, this specific behavioural ecology of briefly interacting with many freshly dehiscent species, but swiftly abandoning them in favor of newly discovered food sources, limits their ability to accurately characterize the entire phenological cycle of flowering plants. However, while honey bee collected pollen is not clearly useful for delimitating the duration or end of flowering, it still provides extremely predicable data indicating the onset of anthesis (Fig. 2) for a wide range of plants (Fig. 4) which is a key parameter that is predicted to be impacted by climate change. Using the on-set of anthesis alone could be a useful indicator of community-level phenological structure, particularly when looking at taxonomic clustering of flowering behaviour (Fig. 8).

Many species inventoried in the field did not occur in our pollen metabarcoding datasets, suggesting that the discriminatory behaviour of foraging bees should be a primary consideration when employing this approach. Despite the obvious preferences displayed by our study colonies, across the 15-week monitoring program our multi-locus pollen metabarcoding approach detected 222 unique plant genera (Fig. 3). Though our ability to monitor these trends is clearly limited by dietary preferences, the generalist foraging behaviour and wide dietary breadth (Wizenberg et al., 2023b) of honey bees demonstrates their ability to rapidly characterize many key players within flowering plant communities (e.g. *Acer, Abies*, *Brassica, Malus, Prunus, Taraxacum, Salix*). In particular, perennial woody taxa at our urban site were well represented within our pollen metabarcoding dataset, suggesting that monitoring the phenology of urban trees is widely feasible. Increasing the number of colonies used for monitoring directly increased our species detection rates, as did employing a multi-locus metabarcoding approach (Fig. 3). Evidently, the establishment of a robust monitoring program for documenting phenological changes within flowering plant communities via the plant-pollinator interface requires the use of multiple large colonies to maximize species detection rates (Fig. 3) and pooled pollen yields (Fig. 5). Interestingly, we did not document a notable asymptote of species detection rates – suggesting that increasing the colonies used to monitor beyond 5 could yield even greater detection rates. Though single-locus characterization is possible, either via *ITS2* which boasts greater taxonomic resolution, or *rbcL* which boasts greater quantitative power, a multi-locus approach is demonstrably superior at detecting a greater number of plant genera (Fig. 3). Here we opted to barcode only to the genus level, as validated in our pilot project (Wizenberg et al., 2023a), however, species-level identification of ASVs for exploring flowering phenology may be attainable through multi-locus identification. Many of the ASVs in our pollen metabarcoding dataset, when barcoded to the species level, did correspond to the species observed at our field site. This suggests that classification of reads at lower taxonomic levels may be achievable if the limitations of this approach are well understood. In particular, the addition of another genetic marker for a tri-locus barcoding program, e.g. *trnL* (Polling et al., 2022), could increase the confidence around species-level taxonomic assignments and should be explored in greater depth.

Employing honey bees to rapidly survey the flowering behaviour of plant communities will revolutionize our ability to monitor these trends on a previously unfathomable scale. Our findings coincide with previous work demonstrating the strength of primary producers as bioindicators of environmental trends (Chowdhury et al., 2023; Desrosiers et al., 2013; Xu & Zhang, 2012). Recent research has suggested that not all invertebrates perform well as bioindicators (Segre et al., 2023), but honey bees have long been used as environmental samplers, gauging the presence of atmospheric pollutants (Kevan, 1999) and metals associated with anthropogenic activity (Skorbiłowicz et al., 2018). Unlike previous work, we are the first to provide evidence that biomonitoring activities could extend beyond plant community composition. Long-term monitoring across multiple spatial scales is key to understanding the stochastic nature of plant phenological structure (Tang et al., 2016; Rafferty et al., 2013), but thus far work has largely been limited to ground-based historical observations, satellite imagery, and digital repeat photography (Fitchett et al., 2015). Utilizing managed honey bee colonies is certainly an innovative approach, and unlike other methodologies, it dually provides valuable data on plant-pollinator interactions. In addition to being dietary generalists, honey bees have several properties that make them highly suitable as bio-monitors of plant phenology: as a highly eusocial species, honey bee colonies have thousands of foragers, thereby increasing their ability to collect pollen over solitary pollinator species. Moreover, as a managed species, honey bees – to our knowledge – have the largest global distribution among the pollinating bees. Their colonies are portable and easily transported, thereby allowing for the development of biomonitoring programs that are standardized and implementable across the world. Hypothetically, standardization across years would be a crucial consideration when establishing a monitoring site, but our work suggests that between colony variation in foraging preferences should be minimally disruptive (Fig. 6). This means that colony turnover, and divergence in colony-level patterns of preferential foraging at different study sites, should not substantially discredit the integrity of a monitoring program.

## CONCLUSION

Here, we present a novel approach for monitoring temporal reshuffling of plant community phenology via honey bee mediated environmental sampling. Our application of DNA metabarcoding to document the timing of pollen release in plant communities represents a substantial improvement over ground-based historical observations and could revolutionize our ability to monitor global change. Metabarcoding-based approaches for studying ecological change have picked up steam in the past decade (Clare et al., 2022; Gold et al., 2021; Sales et al., 2021; Mena et al., 2021; Van Der Heyde et al., 2020; Hallam et al., 2021), as advances in molecular techniques (Soni & Singh, 2022; Ruis et al., 2015; Glenn, 2011) have increased the speed and scale of this type of work. Our work directly builds off of these advances but presents a new avenue for applying environmental DNA approaches to explore evo-eco questions through the lens of a generalist pollinator. Though pollen metabarcoding has widely been used for studying dietary composition and foraging ecology (Leponiemi et al., 2023; Milla et al., 2022; Richardson et al., 2021; Bell et al., 2021), we are the first to demonstrate its power for phenological surveys of plant communities. Hypothetically any pollinating insect could present a reasonable model system for exploring these trends, but the global distribution and generalist foraging behaviour (Quigley et al., 2019; Potts et al., 2010), as well as their wide dietary breadth (Wizenberg et al., 2023b), makes managed honey bee colonies uniquely qualified candidates for standardization across spatial-temporal scales. Establishment of long-term monitoring programs via honey bee collected pollen is dually valuable, as it can indicate changes to phenological structure, and provide fine-scale temporal monitoring of the plant-pollinator interface. Standardized monitoring of these trends will shed light on the cascading impacts of global change and identify divergence consequences of temporal changes in plant-pollinator interactions.

## REFERENCES

Allan, R. P., Barlow, M., Byrne, M. P., Cherchi, A., Douville, H., Fowler, H. J., … & Zolina, O. (2020). Advances in understanding large-scale responses of the water cycle to climate change. Annals of the New York Academy of Sciences, 1472(1), 49–75.

Almas, A. D., & Conway, T. M. (2016). The role of native species in urban forest planning and practice: A case study of Carolinian Canada. Urban Forestry & Urban Greening, 17, 54–62.

Anderson, R., Bayer, P. E., & Edwards, D. (2020). Climate change and the need for agricultural adaptation. Current Opinion in Plant Biology, 56, 197–202.

Bell, K. L., Batchelor, K. L., Bradford, M., McKeown, A., Macdonald, S. L., & Westcott, D. (2021). Optimisation of a pollen DNA metabarcoding method for diet analysis of flying-foxes (Pteropus spp.). Australian Journal of Zoology.

Bell, K. L., De Vere, N., Keller, A., Richardson, R. T., Gous, A., Burgess, K. S., & Brosi, B. J. (2016). Pollen DNA barcoding: current applications and future prospects. Genome, 59(9), 629–640.

Biesmeijer, J. C., & Seeley, T. D. (2005). The use of waggle dance information by honey bees throughout their foraging careers. Behavioral Ecology and Sociobiology, 59, 133–142.

Bonner, C., Sokolov, N. A., Westover, S. E., Ho, M., & Weis, A. E. (2019). Estimating the impact of divergent mating phenology between residents and migrants on the potential for gene flow. Ecology and Evolution, 9(7), 3770–3783.

Callahan, B. J., McMurdie, P. J., Rosen, M. J., Han, A. W., Johnson, A. J. A., & Holmes, S. P. (2016). DADA2: High-resolution sample inference from Illumina amplicon data. Nature Methods, 13(7), 581–583.

Chowdhury, S., Dubey, V. K., Choudhury, S., Das, A., Jeengar, D., Sujatha, B., … & Kumar, V. (2023). Insects as bioindicator: A hidden gem for environmental monitoring. Frontiers in Environmental Science, 11.

Clare, E. L., Economou, C. K., Bennett, F. J., Dyer, C. E., Adams, K., McRobie, B., … & Littlefair, J. E. (2022). Measuring biodiversity from DNA in the air. Current Biology, 32(3), 693–700.

Cooper, E. J., Dullinger, S., & Semenchuk, P. (2011). Late snowmelt delays plant development and results in lower reproductive success in the High Arctic. Plant Science, 180(1), 157–167.

Deiner, K., Bik, H. M., Mächler, E., Seymour, M., Lacoursière-Roussel, A., Altermatt, F., … & Bernatchez, L. (2017). Environmental DNA metabarcoding: Transforming how we survey animal and plant communities. Molecular Ecology, 26(21), 5872–5895.

Desrosiers, C., Leflaive, J., Eulin, A., & Ten-Hage, L. (2013). Bioindicators in marine waters: benthic diatoms as a tool to assess water quality from eutrophic to oligotrophic coastal ecosystems. Ecological Indicators, 32, 25–34.

Elzinga, J. A., Atlan, A., Biere, A., Gigord, L., Weis, A. E., & Bernasconi, G. (2007). Time after time: flowering phenology and biotic interactions. Trends in Ecology & Evolution, 22(8), 432–439.

Fitchett, J. M., Grab, S. W., & Thompson, D. I. (2015). Plant phenology and climate change: Progress in methodological approaches and application. Progress in Physical Geography, 39(4), 460–482

Franks, S. J., & Weis, A. E. (2008). A change in climate causes rapid evolution of multiple life-history traits and their interactions in an annual plant. Journal of Evolutionary Biology, 21(5), 1321–1334.

Franks, S. J., Sim, S., & Weis, A. E. (2007). Rapid evolution of flowering time by an annual plant in response to a climate fluctuation. Proceedings of the National Academy of Sciences, 104(4), 1278–1282.

Franks, S. J., Weber, J. J., & Aitken, S. N. (2014). Evolutionary and plastic responses to climate change in terrestrial plant populations. Evolutionary Applications, 7(1), 123–139.

Gallagher, M. K., & Campbell, D. R. (2020). Pollinator visitation rate and effectiveness vary with flowering phenology. American Journal of Botany, 107(3), 445–455.

Glenn, T. C. (2011). Field guide to next-generation DNA sequencers. Molecular Ecology Resources, 11(5), 759–769.

Gold, Z., Sprague, J., Kushner, D. J., Zerecero Marin, E., & Barber, P. H. (2021). eDNA metabarcoding as a biomonitoring tool for marine protected areas. PLoS One, 16(2), e0238557.

Hallam, J., Clare, E. L., Jones, J. I., & Day, J. J. (2021). Biodiversity assessment across a dynamic riverine system: A comparison of eDNA metabarcoding versus traditional fish surveying methods. Environmental DNA, 3(6), 1247–1266.

Hamann, E., Denney, D., Day, S., Lombardi, E., Jameel, M. I., MacTavish, R., & Anderson, J. T. (2021). Plant eco-evolutionary responses to climate change: Emerging directions. Plant Science, 304, 110737.

Hegland, S. J., Nielsen, A., Lázaro, A., Bjerknes, A. L., & Totland, Ø. (2009). How does climate warming affect plant-pollinator interactions? Ecology letters, 12(2), 184–195.

Inouye, D. W. (2020). Effects of climate change on alpine plants and their pollinators. Annals of the New York Academy of Sciences, 1469(1), 26–37.

Johnson, M. D., Cox, R. D., & Barnes, M. A. (2019). The detection of a non-anemophilous plant species using airborne eDNA. PLoS One, 14(11), e0225262.

Johnson, M. D., Fokar, M., Cox, R. D., & Barnes, M. A. (2021). Airborne environmental DNA metabarcoding detects more diversity, with less sampling effort, than a traditional plant community survey. BMC Ecology and Evolution, 21(1), 1–15.

Keck, F., Blackman, R. C., Bossart, R., Brantschen, J., Couton, M., Hürlemann, S., … & Altermatt, F. (2022). Meta-analysis shows both congruence and complementarity of DNA and eDNA metabarcoding to traditional methods for biological community assessment. Molecular Ecology, 31(6), 1820–1835.

Kevan, P. G. (1999). Pollinators as bioindicators of the state of the environment: species, activity, and diversity. In Invertebrate biodiversity as bioindicators of sustainable landscapes (pp. 373–393). Elsevier

L. Henry, H. Wickham, purrr: Functional Programming Tools (2020).

Lacoursière-Roussel, A., Howland, K., Normandeau, E., Grey, E. K., Archambault, P., Deiner, K., … & Bernatchez, L. (2018). eDNA metabarcoding as a new surveillance approach for coastal Arctic biodiversity. Ecology and Evolution, 8(16), 7763–7777.

Leponiemi, M., Freitak, D., Moreno-Torres, M., Pferschy-Wenzig, E. M., Becker-Scarpitta, A., Tiusanen, M., … & Wirta, H. (2023). Honeybees’ foraging choices for nectar and pollen revealed by DNA metabarcoding. Scientific Reports, 13(1), 14753.

Littlefair, J. E., Zander, A., de Sena Costa, C., & Clare, E. L. (2019). DNA metabarcoding reveals changes in the contents of carnivorous plants along an elevation gradient. Molecular Ecology, 28(2), 281–292.

Lü, J., Wang, R., Sardans, J., Peñuelas, J., Jiang, Y., & Han, X. (2023). An Integrative Review of Drivers and Responses of Grassland Phenology under Global Change. Critical Reviews in Plant Sciences, 1–14.

Mena, J. L., Yagui, H., Tejeda, V., Bonifaz, E., Bellemain, E., Valentini, A., … & Lyet, A. (2021). Environmental DNA metabarcoding as a useful tool for evaluating terrestrial mammal diversity in tropical forests. Ecological Applications, 31(5), e02335.

Milla, L., Schmidt-Lebuhn, A., Bovill, J., & Encinas-Viso, F. (2022). Monitoring of honey bee floral resources with pollen DNA metabarcoding as a complementary tool to vegetation surveys. Ecological Solutions and Evidence, 3(1), e12120.

Morton, E. M., & Rafferty, N. E. (2017). Plant–pollinator interactions under climate change: The use of spatial and temporal transplants. Applications in Plant Sciences, 5(6), 1600133.

Munguía-Rosas, M. A., Ollerton, J., Parra-Tabla, V., & De-Nova, J. A. (2011). Meta-analysis of phenotypic selection on flowering phenology suggests that early flowering plants are favoured. Ecology Letters, 14(5), 511–521.

Parmesan, C. (2006). Ecological and evolutionary responses to recent climate change. Annual Review of Ecology, Evolution, and Systematics, 37, 637–669.

Polling, M., Sin, M., de Weger, L. A., Speksnijder, A. G., Koenders, M. J., de Boer, H., & Gravendeel, B. (2022). DNA metabarcoding using nrITS2 provides highly qualitative and quantitative results for airborne pollen monitoring. Science of the Total Environment, 806, 150468.

Potts, S. G., Biesmeijer, J. C., Kremen, C., Neumann, P., Schweiger, O., & Kunin, W. E. (2010). Global pollinator declines: trends, impacts and drivers. Trends in Ecology & Evolution, 25(6), 345–353.

Qanmber, G., Lu, L., Liu, Z., Yu, D., Zhou, K., Huo, P., … & Yang, Z. (2019). Genome-wide identification of GhAAI genes reveals that GhAAI66 triggers a phase transition to induce early flowering. Journal of Experimental Botany, 70(18), 4721–4736.

Quigley, T. P., Amdam, G. V., & Harwood, G. H. (2019). Honey bees as bioindicators of changing global agricultural landscapes. Current Ppinion in Insect Science, 35, 132–137.

Rafferty, N. E., Diez, J. M., & Bertelsen, C. D. (2020). Changing climate drives divergent and nonlinear shifts in flowering phenology across elevations. Current Biology, 30(3), 432–441.

Rafferty, N. E., & Ives, A. R. (2011). Effects of experimental shifts in flowering phenology on plant–pollinator interactions. Ecology Letters, 14(1), 69–74.

Rafferty, N. E., CaraDonna, P. J., Burkle, L. A., Iler, A. M., & Bronstein, J. L. (2013). Phenological overlap of interacting species in a changing climate: an assessment of available approaches. Ecology and Evolution, 3(9), 3183–3193.

Ratnieks, F. L., & Shackleton, K. (2015). Does the waggle dance help honey bees to forage at greater distances than expected for their body size?. Frontiers in Ecology and Evolution, 3, 31.

Richardson, R. T. (2022). Controlling critical mistag-associated false discoveries in metagenetic data. Methods in Ecology and Evolution, 13(5), 938–944.

Richardson, R. T., Curtis, H. R., Matcham, E. G., Lin, C. H., Suresh, S., Sponsler, D. B., … & Johnson, R. M. (2019). Quantitative multi-locus metabarcoding and waggle dance interpretation reveal honey bee spring foraging patterns in Midwest agroecosystems. Molecular Ecology, 28(3), 686–697.

Richardson, R. T., Eaton, T. D., Lin, C. H., Cherry, G., Johnson, R. M., & Sponsler, D. B. (2021). Application of plant metabarcoding to identify diverse honeybee pollen forage along an urban–agricultural gradient. Molecular Ecology, 30(1), 310–323.

Richardson, R. T., Sponsler, D. B., McMinn-Sauder, H., & Johnson, R. M. (2020). MetaCurator: A hidden Markov model-based toolkit for extracting and curating sequences from taxonomically-informative genetic markers. Methods in Ecology and Evolution, 11(1), 181–186.

Rius, M., Bourne, S., Hornsby, H. G., & Chapman, M. A. (2015). Applications of next-generation sequencing to the study of biological invasions. Current Zoology, 61(3), 488–504.

Rodríguez-Pérez, J., & Traveset, A. (2016). Effects of flowering phenology and synchrony on the reproductive success of a long-flowering shrub. AoB Plants, 8.

Ruppert, K. M., Kline, R. J., & Rahman, M. S. (2019). Past, present, and future perspectives of environmental DNA (eDNA) metabarcoding: A systematic review in methods, monitoring, and applications of global eDNA. Global Ecology and Conservation, 17, e00547.

Rymer, P. D., Johnson, S. D., & Savolainen, V. (2010). Pollinator behaviour and plant speciation: can assortative mating and disruptive selection maintain distinct floral morphs in sympatry?. New Phytologist, 188(2), 426–436.

Sales, N. G., Wangensteen, O. S., Carvalho, D. C., Deiner, K., Præbel, K., Coscia, I., … & Mariani, S. (2021). Space-time dynamics in monitoring neotropical fish communities using eDNA metabarcoding. Science of the Total Environment, 754, 142096.

Savage, J. A. (2019). A temporal shift in resource allocation facilitates flowering before leaf out and spring vessel maturation in precocious species. American Journal of Botany, 106(1), 113–122.

Segre, H., Kleijn, D., Bartomeus, I., WallisDeVries, M. F., de Jong, M., van der Schee, M. F., … & Fijen, T. P. (2023). Butterflies are not a robust bioindicator for assessing pollinator communities, but floral resources offer a promising way forward. Ecological Indicators, 154, 110842.

Shackleton, K., Balfour, N. J., Al Toufailia, H., James, E., & Ratnieks, F. L. (2023). Honey bee waggle dances facilitate shorter foraging distances and increased foraging aggregation. Animal Behaviour, 198, 11–19.

Shen, M., Piao, S., Dorji, T., Liu, Q., Cong, N., Chen, X., … & Zhang, G. (2015). Plant phenological responses to climate change on the Tibetan Plateau: research status and challenges. National Science Review, 2(4), 454–467.

Skorbiłowicz, E., Skorbiłowicz, M., & Cieśluk, I. (2018). Bees as bioindicators of environmental pollution with metals in an urban area. Journal of Ecological Engineering, 19(3), 229–234.

Soni, H. K., & Singh, A. (2022). Application of Next-Generation Sequencing in Plant Molecular Ecology. In Plant Ecogenomics (pp. 47–81). Apple Academic Press.

Tang, J., Körner, C., Muraoka, H., Piao, S., Shen, M., Thackeray, S. J., & Yang, X. (2016). Emerging opportunities and challenges in phenology: a review. Ecosphere, 7(8), e01436.

Tereshko, V., & Lee, T. (2002). How information-mapping patterns determine foraging behaviour of a honey bee colony. Open Systems & Information Dynamics, 9(2), 181–193.

Tun, W., Yoon, J., Jeon, J. S., & An, G. (2021). Influence of climate change on flowering time. Journal of Plant Biology, 64(3), 193–203.

Van Der Heyde, M., Bunce, M., Wardell-Johnson, G., Fernandes, K., White, N. E., & Nevill, P. (2020). Testing multiple substrates for terrestrial biodiversity monitoring using environmental DNA metabarcoding. Molecular Ecology Resources, 20(3), 732–745.

Vasiliev, D., & Greenwood, S. (2021). The role of climate change in pollinator decline across the Northern Hemisphere is underestimated. Science of the Total Environment, 775, 145788.

Wadgymar, S. M., & Weis, A. E. (2017). Phenological mismatch and the effectiveness of assisted gene flow. Conservation Biology, 31(3), 547–558.

Weis, A. E., & Kossler, T. M. (2004). Genetic variation in flowering time induces phenological assortative mating: quantitative genetic methods applied to Brassica rapa. American Journal of Botany, 91(6), 825–836.

Weis, A. E., Winterer, J., Vacher, C., Kossler, T. M., Young, C. A., & LeBuhn, G. L. (2005). Phenological assortative mating in flowering plants: the nature and consequences of its frequency dependence. Evolutionary Ecology Research, 7(2), 161–181.

Wellmer, F., & Riechmann, J. L. (2010). Gene networks controlling the initiation of flower development. Trends in Genetics, 26(12), 519–527.

Wickham, H. (2016). ggplot2: elegant graphics for data analysis. R package version 3.4.2.

Wilke, C. O. (2018). Ggridges: Ridgeline plots in’ggplot2’. R package version 0.5.1.

Wizenberg, S. B., Newburn, L. R., Pepinelli, M., Conflitti, I. M., Richardson, R. T., Hoover, S. E., … & Zayed, A. (2023a). Validating a multi-locus metabarcoding approach for characterizing mixed-pollen samples. Plant Methods, 19(1), 120.

Wizenberg, S. B., Newburn, L. R., Richardson, R. T., Pepinelli, M., Conflitti, I. M., Moubony, M., … & Zayed, A. (2023b). Environmental metagenetics unveil novel plant-pollinator interactions. Ecology and Evolution, 13(11), e10645.

Xie, S. P., Deser, C., Vecchi, G. A., Collins, M., Delworth, T. L., Hall, A., … & Watanabe, M. (2015). Towards predictive understanding of regional climate change. Nature Climate Change, 5(10), 921–930.

Xu, J., & Zhang, M. (2012). Primary consumers as bioindicator of nitrogen pollution in lake planktonic and benthic food webs. Ecological Indicators, 14(1), 189–196.

Ziska, L. H., & Beggs, P. J. (2012). Anthropogenic climate change and allergen exposure: the role of plant biology. Journal of Allergy and Clinical Immunology, 129(1), 27–32.

